# Designer protein assemblies with tunable phase diagrams in living cells

**DOI:** 10.1101/2020.06.03.131433

**Authors:** Meta Heidenreich, Joseph M. Georgeson, Emanuele Locatelli, Lorenzo Rovigatti, Saroj Kumar Nandi, Avital Steinberg, Yotam Nadav, Eyal Shimoni, Samuel A. Safran, Jonathan P. K. Doye, Emmanuel D. Levy

## Abstract

The self-organization of proteins into specific assemblies is a hallmark of biological systems. Principles governing protein-protein interactions have long been known. However, principles by which such nanoscale interactions generate diverse phenotypes of mesoscale assemblies, including phase-separated compartments, remains challenging to characterize and understand. To illuminate such principles, we create a system of two proteins designed to interact and form mesh-like assemblies in living cells. We devise a novel strategy to map high-resolution phase diagrams *in vivo*, which provide mesoscale self-assembly signatures of our system. The structural modularity of the two protein components allows straightforward modification of their molecular properties, enabling us to characterize how point mutations that change their interaction affinity impact the phase diagram and material state of the assemblies *in vivo*. Both, the phase diagrams and their dependence on interaction affinity were captured by theory and simulations, including out-of-equilibrium effects seen in growing cells. Applying our system to interrogate biological mechanisms of self-assembly, we find that co-translational protein binding suffices to recruit an mRNA to the designed micron-scale structures.

## Introduction

The self-organization and proper function of complex systems involve elaborate spatiotemporal coordination of their constituent elements. Cells organize their contents into organelles, which have been classically viewed as membrane-bound structures. However, in recent years, an increasing number of studies describe a fundamentally different type of organelles that form by phase separation and are not membrane-bound. These organelles, also called biomolecular condensates, include P-Bodies^1^, stress granules^2^, nucleoli^3^ among many others^4–6^. The functions associated with these organelles are diverse^5,7,8^, ranging from pre-mRNA processing^9^ and translation regulation^10,11^, to signalling^12^, or to the formation of eye lenses^13^. The increasingly frequent discovery of such organelles reflects that we are only beginning to grasp the complexity underlying the proteome’s spatial organization and begs for a molecular understanding of the process of phase separation in living cells.

In phase separation, thousands of copies of identical molecules cluster and interact together, implying that small changes in molecular properties of components, e.g., by mutation, can propagate and dramatically impact macroscopic phenotypes of assembly^14^. For example, mutations increasing the viscosity of FUS and Huntington exon 1 condensates have been associated with debilitating diseases such as amyotrophic lateral sclerosis (ALS), frontotemporal dementia (FTD)^15,16^, and Huntington^17,18^. However, there is little understanding of how these mutations act at the molecular level to change the phase behaviour and viscosity of condensates. In order to bridge this gap, it is crucial to connect biophysical properties of proteins to mesoscale phenotypes of their assembly inside of living cells.

Establishing such a nanoscale-mesoscale connection with natural condensates is hardly possible due to their compositional and regulatory complexity. Creating synthetic condensates offers a powerful alternative, as both the structure and biophysical properties of the components can be known by design. Furthermore, if the proteins employed are orthogonal to the living system, no active cellular regulation is expected to take place. Previous work based on synthetic proteins showed that increasing multivalence of the components promotes their phase separation^19,20^, and revealed how distinct client proteins can be differentially recruited to condensates^4^. However, detailed molecular modeling of these systems is impossible, since the interaction affinity between individual components was fixed^19^ or unknown^20^, and the contribution of intra-*versus* inter-molecular interactions was also unknown. This prompted us to design a synthetic system providing control over these nanoscale properties. Equally important, we set out that such a system should allow the direct visualization of mesoscale self-assembly signatures, through high-resolution phase diagrams measured in living cells.

## Results

### A synthetic two-protein system that forms condensates in vivo

A quantitative and detailed molecular understanding of biophysical and biological mechanisms of mesoscale self-assembly requires a system where all parameters, namely the components, their structure, and their physical interactions, are known. To this aim, we developed a synthetic system in which these properties are controlled by design. The system comprises two protein components that interact with affinities tunable by point mutation. Each component is designed in a modular fashion and consists of three structured domains linked by short flexible linkers. As we know from previous work that multivalence is a critical property of molecules undergoing phase separation^19,21^, both components are multivalent. The first component contains a homo-dimerization domain, a red fluorescent protein (RFP), and the protein Im2. The second component contains a homo-tetramerization domain, a yellow fluorescent protein (YFP), and the protein E9, which interacts specifically with Im2 (Figure 1a, Methods, Table S1). Importantly, unlike in other synthetic systems^19,20^, intramolecular interactions are restricted by an incompatibility between the distances separating the termini to which interaction domains are fused, equal to 18 nm on the dimer and only 4 nm on the tetramer (Figure 1a, b).

**Figure 1.**
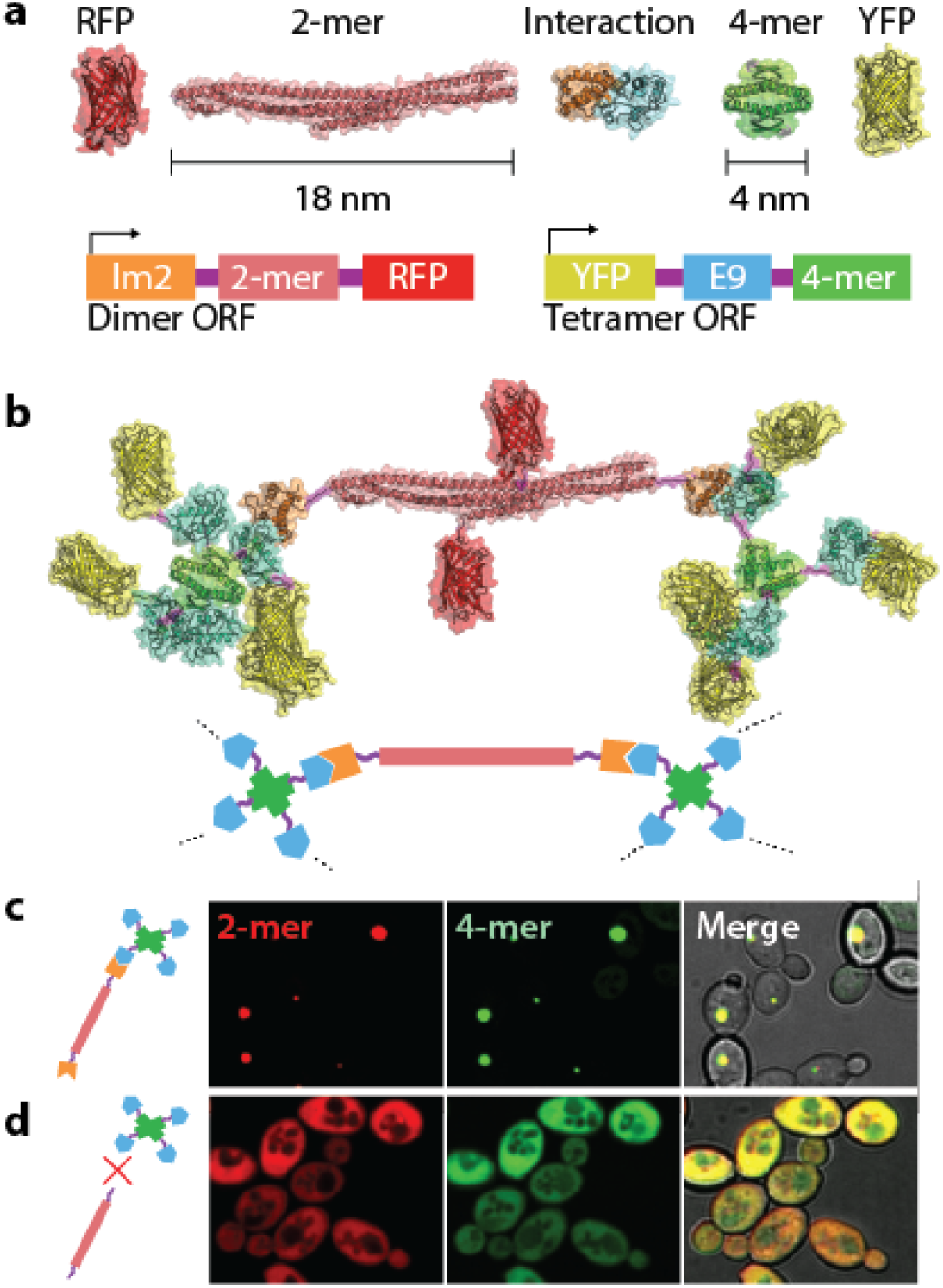
A synthetic system for controlled phase separation in living cells. **a.** The components, each encoded in one ORF, consist of three domains connected by flexible linkers: An interaction domain, an oligomerization domain and a fluorescent protein. The Colicin (E9, cyan) and Immunity (Im2, orange) proteins serve as interaction modules, where affinity is controllable by mutation. A dimer and tetramer of known structure (Table S1) served as divalent and tetravalent scaffolds. We fused Im2 and a red fluorescent protein (RFP) to the dimer, and E9 and a yellow fluorescent protein (YFP) to the tetrameric scaffold. **b**. Illustrative structure of a dimer interacting with two tetramers, and cartoon representation underneath. **c**. The system undergoes self-assembly and forms punctate structures in living yeast cells. **d**. In absence of the Im2 interaction module no punctate structure is formed.

### The system phase separates in vivo

We co-expressed the dimer and tetramer components in yeast cells. Using fluorescence microscopy, we observed the formation of sub-micron to micron-scale punctate assemblies where the tetramer and dimer co-localized (Figure 1c, Movie 1), suggesting that the system undergoes phase separation and forms condensates. The assembly of this system was dependent on the specific interaction between E9 and Im2, as condensates were neither observed when co-expressing the tetramer with a dimer lacking the Im2 domain (Figure 1d), nor were they observed in haploid cells that expressed only one of the components (Figure S1). The assemblies were not membrane-bound, as visualized by electron microscopy (Figure S2).

### Revealing phase diagrams in vivo at high-resolution

In a single cell, phase separation of the dimer-tetramer mixture generates two protein phases: one is dense and corresponds to the condensate, the other is dilute and corresponds to freely diffusing components in the cytoplasm (Figure 2a)^19,21–24^. The conditions under which phase separation of a system occurs at equilibrium is described by its phase diagram, with the binodal defining phase boundaries. We developed a lattice model of interacting dimers and tetramers (Figure 2b, Text S1) to model the phase diagram of our system (Figure 2c). Dimer and tetramer concentrations outside of the binodal do not drive phase separation, either because the concentration of components is too low relative to their interaction affinity (Figure 2d), or because imbalance in the components’ stoichiometry inhibits the propagation of their interactions (Figure 2e). Interestingly, cells without condensates have not undergone phase separation and should fall outside of the binodal. Thus, the region of concentrations that is absent in these cells should reveal the phase boundary of this system (Figure 2f). Such an approach offers the unique opportunity to map a high-resolution phase diagram *in vivo*, because the phase-space can be defined along two continuous coordinates corresponding to the concentrations of each of the components. Unlike temperature or pressure, protein concentration can be tuned over several orders of magnitude and can be measured readily from fluorescence intensity across thousands of single cells.

**Figure 2.**
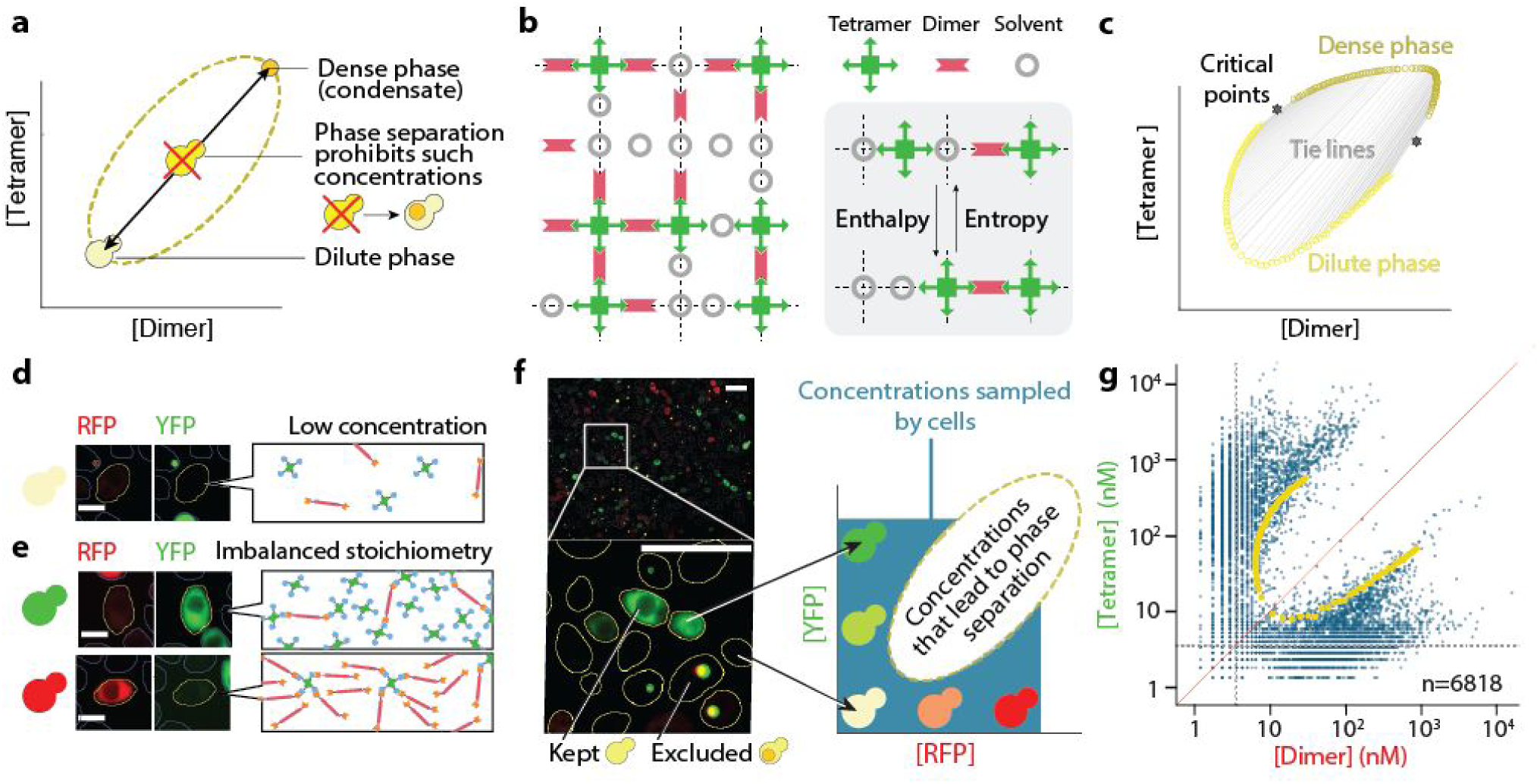
Characterizing phase diagrams in living cells. **a.** Phase separation is the de-mixing of a solution into a high- and a low-concentration phase. The phase diagram describes when the system phase separates in a given parameter space, here defined by the concentrations of two interacting components. Concentrations within the binodal (yellow dotted line) are not stable and will not be observed at equilibrium. The crossed-out cell shows unstable concentrations leading to demixing into low- and high-concentration phases. **b**. A minimal lattice model captures the essence of phase separation, whereby the chemical potential of the dimer and tetramer exhibit two minima (Text S1), the first with high entropy and low enthalpy (dilute phase), and the second with low entropy and high enthalpy from the bonding energy (dense phase). **c.** Based on this lattice model we derive a phase diagram showing the binodal, two critical points and ties lines. **d.** Cells without condensate may have concentrations of both components that are too low. **e**. Alternatively, cells without condensate may exhibit an imbalanced stoichiometry, where binding sites of the component of lower concentration are saturated with the component in excess. **f.** Cells are imaged, segmented, and cells with condensates are excluded. The concentrations of dimer (RFP, red), and tetramer (YFP, green) binding sites are recorded and plotted against each other. Both components are co-expressed stochastically, so each cell samples one point of the phase diagram. Scale bar: 10 µm **g.** *In vivo* phase diagram of our synthetic system containing wild-type Im2 and E9 interacting with a reported affinity of 15 nM (Table 1). Each point corresponds to the concentrations of dimer (x-axis), and tetramer subunits (y-axis) in a single cell. The scatterplot shows data from 6818 cells. The red line highlights the diagonal. Grey dotted lines delimit background fluorescence levels below which concentrations cannot be estimated reliably (∼3.5 nM). The yellow points show an overlay of the binodal computed based on the lattice model (Text S1).

To characterize such a phase diagram, we thus created yeast strains co-expressing the dimer and tetramer components independently, such that each cell sampled a different point of the phase space. We imaged thousands of single cells and estimated the components’ concentrations from fluorescence intensity (Figure S3), excluding cells containing a condensate to ascertain reliable concentration measurements (Figure 2f, Methods). As predicted, the density distribution of cells revealed the phase boundary of the system (Figure 2g).

**Table 1.** Im2 variants previously reported^25^ and used to modulate the interaction affinity between the dimer and tetramer.

| Im2 mutation | $K_d$ with E9 (M) |
| --- | --- |
| D33L | $4.8 \pm 0.3 \times 10^{-11}$ |
| WT | $1.5 \pm 0.1 \times 10^{-8}$ |
| E30A | $2.8 \pm 1.6 \times 10^{-7}$ |
| P56A | $2.1 \pm 0.7 \times 10^{-6}$ |
| V37A | $9.3 \pm 4.4 \times 10^{-6}$ |

### Modeling the phase diagram measured in vivo

The phase boundary appears as an area where cell density approaches zero. The scarcity of cells sampling concentrations beyond 10 µM prevented visualizing closed boundaries, giving rise to a half-ellipsoid. We modelled the expected boundaries using a minimal lattice model (Figure 2b,c and Text S1), as well as a thermodynamic perturbation theory developed for patchy particles matching the geometry of our proteins (Figure S4, Text S2). Both methodological approaches recapitulated our observations: the half-ellipsoid aligns along the diagonal where the stoichiometry of both components’ binding sites is equal (Figure S5). Indeed, a balanced stoichiometry gives rise to a lower energy assembly, where enthalpy is maximal with all binding sites satisfied, thus favoring phase separation. As stoichiometries become unbalanced (e.g., 1:10 or 10:1), the component present in excess saturates all binding sites of its partner, which inhibits propagation of interactions and phase separation (Figure 2e).

### Tuning the phase diagram and effective viscosity of condensates with interaction affinity

The nature of the interaction domains used in this system allows both lowering and increasing the affinity by single point mutations^25^. We created four new variants for the dimer, which contained point mutations modulating the dissociation constant between Im2 and E9 domains across five orders of magnitude, from 10^−11^ to 10^−6^ M (T able 1).

We imaged yeast cells co-expressing the tetramer with the new dimer variants, and generated their *in vivo* phase diagrams (Figure 3 and Figure S6). Mutants interacting with an affinity lower than that of the wild-type domains showed a shift in their phase diagram. The half-ellipsoid underwent a translation along the diagonal, towards higher concentrations. Such a translation was expected, as lower interaction affinities require higher concentrations for binding. The same effect is reproduced with the two theoretical approaches we put forward (Figure S5). Interestingly, the mutant with an affinity higher than the wild type (48 pM) revealed a complex behavior: the minimal concentration of tetramer required for phase separation increased, as reflected in the upward shift of the phase boundary (yellow region, Figure 3a). Such increase suggested the Im2 D33L mutant was structurally impaired and did not interact with the reported interaction affinity.

**Figure 3.**
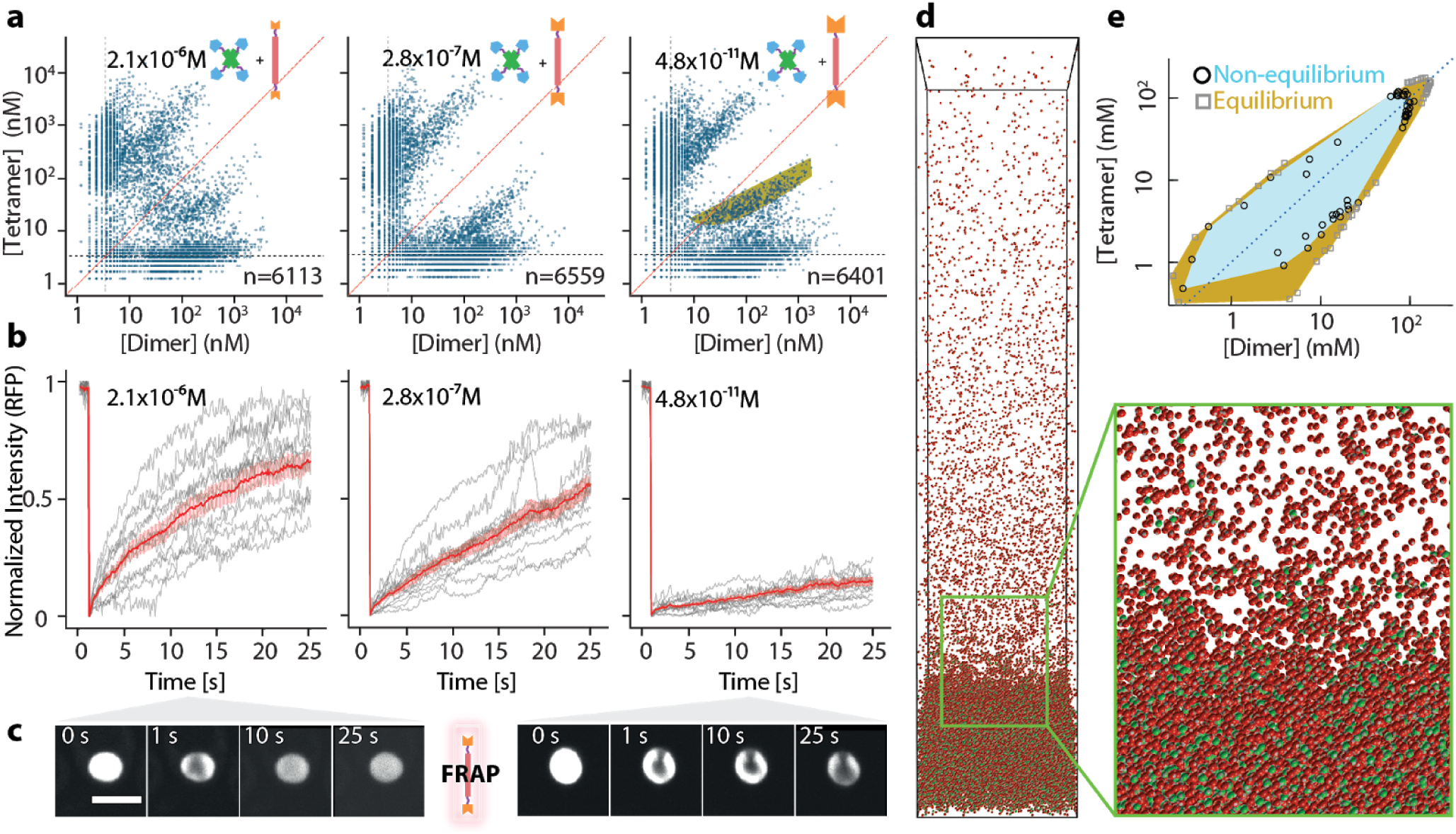
Influence of affinity on *in vivo* phase separation. **a.** Phase diagrams of the tetramer with the dimer carrying three different affinities, as indicated. The red line highlights the diagonal. The grey dotted lines indicate the fluorescence accuracy limit (∼3.5 nM), below which autofluorescence increases. The yellow band highlights a region where phase separation occurs with wild type Im2, but does not with the high affinity mutant. **b**. FRAP experiments were carried out for three pairs of components varying in their interaction affinity. Increasing the interaction affinity increased the effective viscosity of the condensate, n=14. **c.** Example of two condensates recovering after photobleaching. Low affinity interaction (left) shows faster recovery when compared to condensates involving higher affinities (right). Scale bar: 5 µm. **d**. Sedimentation molecular dynamics simulation of patchy particles. Several simulations were conducted at equilibrium or out-of-equilibrium while sampling different concentrations of dimer and tetramer. The protein osmotic pressure as a function of density was inferred from each simulation and used to evaluate the phase boundaries (Figure S8, S9). **e**. The phase diagram of the patchy mixture computed with equilibrium and non-equilibrium simulations (squares and circles, respectively).

This discrepancy led us to examine diffusion dynamics of components within condensates. Fast diffusion requires components to be unbound, and their probability to exist in the unbound state is inversely proportional to their interaction affinity (Text S3). Thus, we expect high affinity interactions to yield condensates with slow diffusion dynamics, whereas lower affinities should yield faster diffusion dynamics. To test this hypothesis, we measured fluorescence recovery after photobleaching (FRAP) of the condensates. Considering low, medium and high affinity interactions (2.1e10^−6^ M, 2.8e10^−7^ M, and 4.8e^-11^ M), the mean fluorescence recovery after 25 seconds reached 65±4%, 56±4% and 15±2%, respectively (Figure 3b and c, Figure S6, Movie 2). Individual traces show pronounced variability in the recovery profiles, especially at low affinities, which might reflect differences in condensate density as well as differences in the fraction of bonded components (Figure S7). On average however, higher interaction affinity led to slower diffusion of components, consistent with the effective viscosity of the condensates being controlled by interaction affinity. Importantly, the slower recovery of the D33L Im2 mutant implies that it does interact with a higher affinity than wild-type Im2, which is in conflict with the observed shrinkage in phase boundaries (yellow region, Figure 3a). We hypothesized that this apparent contradiction originated in kinetics. At high affinity, the kinetics of unbinding events is very slow, which can trap the system in states where both components have a non-optimal distribution of bonds in the network. Nonetheless, dimers need to be completely bonded to mediate cluster growth, whereas tetramers require only two out of four bonds to mediate such growth. Consequently, misplaced bonds in a tetramer-poor system would hinder the formation of a network more than they would in a tetramer-rich system. This idea led us to compare the regions where phase separation occurs in equilibrium versus out-of-equilibrium molecular dynamics simulations of patchy particles (Figure 3d). These simulations confirmed the picture sketched above by revealing a shift in the lower branch of the phase diagram, while the upper branch remained essentially unmoved (Figure 3e, Figure S8, S9, Text S2.4).

### Cotranslational assembly suffices to direct mRNA subcellular localization

The spatial organization of translation is achieved by mRNA trafficking and localization^26,27^. Trafficking involves molecular motors that transport mRNAs across micrometers in yeast cells, and up to tens of centimeters in neurons^26^. After reaching their target site, mRNAs need to be anchored in order to not diffuse away during translation. Interestingly, anchoring could be achieved by the proteins being synthesized, if they bind localized partners co-translationally. This mechanism had, in fact, been suggested to mediate the localization of mRNAs encoding myosin heavy chain in developing cultured skeletal muscles^28^.

However, considering such a biological system, it is hard to address whether cotranslational assembly could be sufficient to anchor mRNAs at a target location, given that alternative or complementary mechanisms could be at play. Uniquely, our synthetic system makes it possible to address this question directly because we know that its components have neither evolved to bind their own mRNA, nor RNAs in general. We fused the mRNA encoding the dimer component to a sequence enabling its tracking in live cells^29^. In these experiments, we used a tetramer component fused to a blue fluorescent reporter, so that green fluorescence was solely reporting on mRNA localization. Live cell imaging revealed that mRNAs diffused throughout the cell and attached to the condensate when they encountered it. Surprisingly, multiple mRNAs could co-localize and appeared to nucleate the formation of the condensate (Figure 4a, Movie 3). In contrast, an mRNA coding for a protein that does not bind to the condensate did not co-localize with it (F igure 4b, Movie 4). Finally, treatment of cells with puromycin, a drug that dissociates ribosomes from mRNA, released the dimers’ mRNA from the condensate within minutes (Figure 4c, Movie 5). These results point to cotranslational assembly being a direct and sufficient driver of localization for the mRNA encoding the dimer.

**Figure 4.**
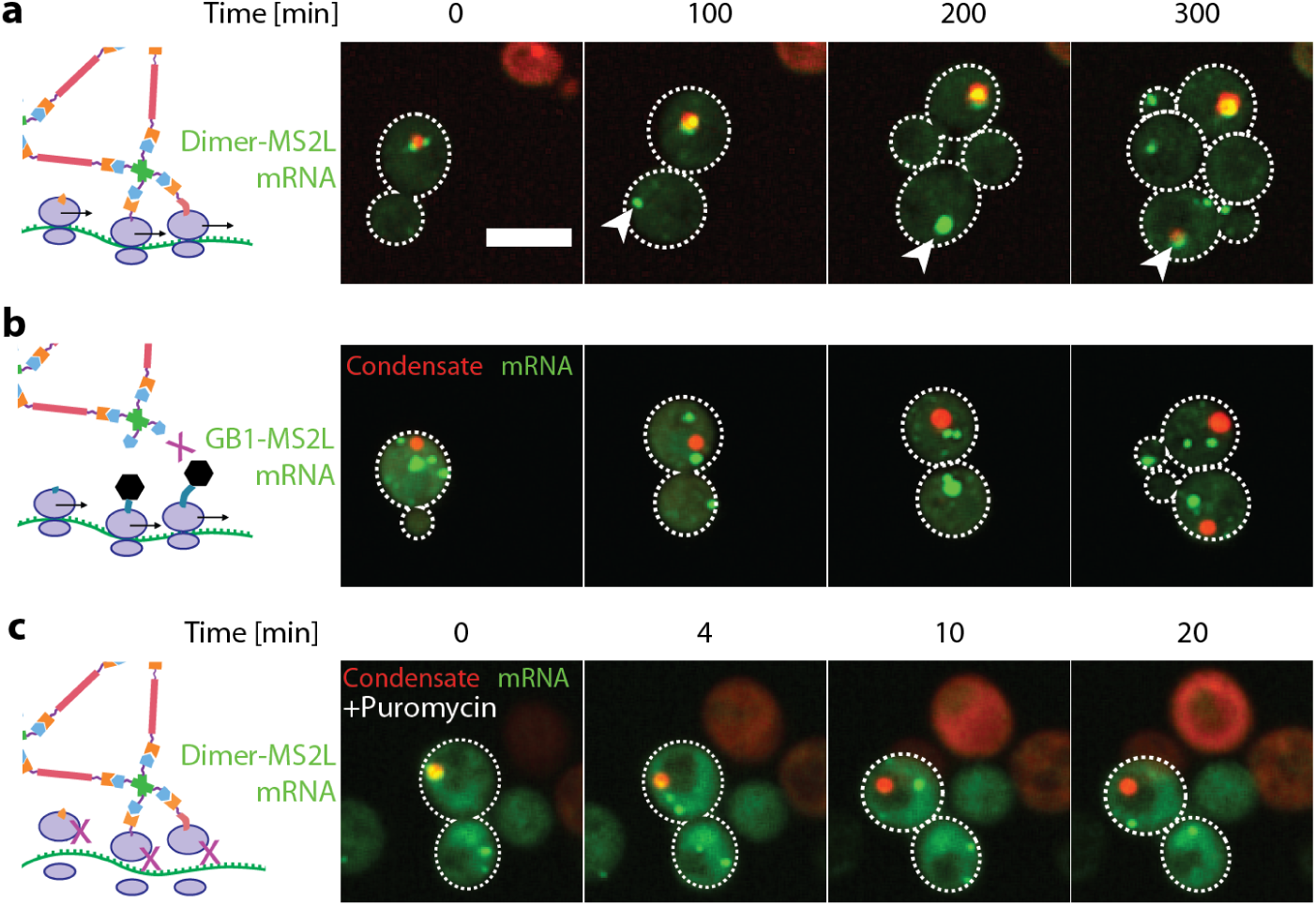
Cotranslational assembly recruits mRNA to the condensate. **a.** The mRNA of the dimeric component was tagged with the MS2 sequence, and appears in live cells as green fluorescent puncta^29^. The tetrameric component did not contain YFP, so the condensates are shown with red fluorescence only. The mRNA molecules encoding for the dimer co-localize with the condensate. **b.** mRNAs of a control protein (GB1) that does not bind to the condensate do not colocalize with it. **c.** Puromycin treatment dissociates ribosomes from mRNA and releases the dimers’ mRNA from the condensate. Scale bar: 5 µm.

## Discussion and conclusions

The system designed in this work presents a number of unique features. Notably, the folded nature of interaction domains, together with the defined geometry of oligomerization domains provide unprecedented control over the biophysical and structural properties of the components. At the same time, we introduce a novel strategy using single cells as individual “test-tubes” to map high-resolution phase diagrams *in vivo*, in a direct and simple way. Combined, these properties create a powerful experimental system to relate nanoscale to mesoscale phenotypes of self-assembly from first principles. We explore this relationship by characterizing how mutations changing the interaction affinity between the two components impact their phase behavior. Interestingly, numerous additional parameters such as linker properties, electrostatics, or valence could be tuned independently from one another, and their impact on phase separation characterized and modelled in the same way.

The system does not only allow addressing biophysical questions with the use of *in vivo* phase diagrams. It is also well suited to explore synergism between mesoscale protein self-assembly and the cell. Recent works have revealed cotranslational assembly of complexes as a widespread mechanism^30,31^ actively shaped by evolution^32,33^. Several mechanisms for mediating interactions between RNA and proteins in condensates are known^34–36^, and our results suggest cotranslational assembly as a new such mechanism.

The design of mesoscale synthetic protein assemblies is becoming increasingly powerful to create new materials^37–39^ and functions^40–42^. Moreover, as we are only beginning to grasp the complexity of proteome self-organization, new approaches are needed for characterizing and understanding mesoscale properties of protein self-assembly in cells^43–48^. In this context, our synthetic condensate will serve to evaluate and calibrate physical models of self-assembly *in vivo*, form a basis for developing new biomaterials and scaffolds in living cells, and constitute powerful tools to interrogate biological mechanisms of protein assembly.

## Methods

### Design

The synthetic system introduced in this work relies on homo-oligomerization to create multivalent components. We chose specific homo-oligomerization domains so as to avoid intra-molecular interactions between components. Specifically, we selected a large dimer and a small tetramerization domain such that the dimers could bridge across two tetramers, but could not bind two sites on the same tetramer. The dimer (PDB code: 4LTB, the dimerization domain of TRIM25^49^), consists of an antiparallel coiled-coil, where both N-termini are 18 nm apart. The tetramer is comparatively small and corresponds to the tetramerization domain of p53 (PDB code: 1AIE^50^).

To avoid non-specific interactions of the dimer protein we mutated highly exposed and hydrophobic surface residues to charged ones (Y22D, I92D). For the tetrameric component, we used the wild-type sequence of the tetramerization domain of human p53, from amino acid 326 to 356. The yellow fluorescent reporter Venus bearing the A206K mutation^51^ was fused to the tetramer, and FusionRed^52^ to the dimer. Both are monomeric to prohibit unspecific interactions between the components. The interaction domains were derived from the bacterial toxin - antitoxin system E9/Im2, where different affinities were introduced by point mutations of Im2 (Table 1). A H103A mutant of E9 was used to inhibit its toxic DNAse activity. Upon initial expression in yeast cells, the dimer component showed a tendency for nuclear localization. We thus fused a nuclear export signal (NES) LAEKLAGLDIN^53^ at its N-terminus, which led to its cytosolic localization.

### Plasmids and Strains

To achieve stochastic expression of each component in yeast cells, each ORF was inserted into a separate plasmid. The tRNA adaptation index of sequences for all components was optimized for *S. cerevisiae*. Designed sequences were inserted into American Type Culture Collection (ATCC) yeast cassettes^54^ using the Polymerase Incomplete Primer Extension (PIPE) cloning method^55^. For stoichiometric expression in Movie 1, sequences were inserted into M3925 plasmids^56^ for genomic integration. Both components were cloned downstream of the yeast TDH3 promoter. The selection markers for the dimer and tetramer were hygromycin and G418, respectively. Cloning was performed in *E. coli* DH5α cell. Plasmids were subsequently isolated, verified by sequencing, and transformed into BY4741 (tetramer) or BY4742 (dimer) strains of S288C^57^. Expression in haploid cells was verified by microscopy and yeast were subsequently mated, creating diploid cells containing both plasmids. For investigating the localization of mRNA, a modified version of the mTAG method^29^ was used. Instead of inserting the MS2 loops to the 3’UTR by using the Cre-Lox system, we used CRISPR/Cas9. We used the plasmid bRA89^58^, which carries both, the ORF for Cas9, and the guide RNA. The guide RNA was designed using CRISPR-ERA^59^, to target the TRP2 locus (GTGGACAATCTCACCAGCGT) and the dimer with the wild type Im2, including the MS2 loops in its 3’UTR, was inserted. For the insertion cassette, three pieces were amplified: one from the promoter to the stop codon, one from the stop codon to the end of UTR containing 12 MS2 loop repeats, and one from the end of the 3’ UTR to the end of the terminator. The primers for this amplification contained 40 bp homology regions to the TRP2 locus on the flanking regions, and to each other in overlapping regions. The PCR products were treated with DPN1 (New England Biolabs inc.) and purified using the Agencourt AMPure XP system. We transformed 20 µl of competent BY4742 cells with 1 µl (1 µg/µl) pRA89 (TRP2) and 200-300 ng of each module of the insertion cassette. After inserting the dimer, cells were co-transformed with the plasmid carrying CP-3xGFP and a plasmid carrying the tetramer fused to BFP, instead of Venus. For the negative control, the insertion cassette was comprised of three fragments: one with the TDH3 promoter and GB1, one with the MS2L containing 3’ UTR and one with the CYC terminator. The three fragments were purified with the Agencourt AMPure XP system, joined by PCR, and the resulting piece was again purified. 500 ng of the product was co-transformed with 1 µg of pRA89 (TRP2) to 20 µl competent BY4742 cells. The resulting strain was co-transformed with the CP-3xGFP plasmid as well as a single plasmid carrying both components. Finally, all strains were verified by sequencing. We note that one of the 12 MS2 loops was missing in the negative control.

### Microscopy and Image Processing

Cells were imaged with an Olympus IX83 microscope coupled to a Yokogawa CSU-W1 spinning disc confocal scanner with dual Hamamatsu ORCA-Flash4.0 V2 sCMOS cameras. 16-bit images were acquired for Brightfield and two confocal illumination schemes: GFP channel (Ex 488 nm, Toptica 100 mW | Em 525/50 nm, Chroma ET/m), and RFP channel (Ex 561 nm, Obis 75 mW | Em 609/54 nm, Chroma ET/m). Imaging was performed with a 60x, 1.35 NA, oil immersion objective (UPLSAPO60XO, Olympus) and FRAP experiments were carried out with a 100x, 1.4 NA, oil-immersion objective (UPLSAPO100XO, Olympus). Automated imaging was performed with a motorized XY stage, onto which a piezo-stage (Mad City Labs) was mounted and used for acquiring z-stacks. For phase diagrams, we acquired seven z-stack images for each fluorescent channel, and the average intensity projection was used. For time lapse series, eight z-stacks were acquired, and the maximum intensity projection was used.

### Sample preparation for imaging

A liquid handling robot (Tecan Evo 200) was used to prepare Greiner™ 384-well glass-bottom optical imaging plates. For imaging, 0.5 µl of saturated cell suspension was transferred into an optical plate with SD medium and grown for 6 h to logarithmic growth. For time lapse series, cells were grown to an OD600 of 0.4-0.8, transferred to matrical 96-well glass-bottom plates, and covered with 0.5% Agarose/SD containing the respective resistance marker. For time lapse series of puromycin treatment, cells were not covered with agarose and puromycin was added to the cells after 6 minutes of imaging, to a final concentration of 62 mM. For FRAP experiments, cells were grown and let at saturation for two weeks to generate large condensates. Cells were subsequently fixed with ConA in an optical 96-well plate, as previously described^59,60^, and FRAP experiments were carried out 6 h after their inoculation into fresh media.

### Image analysis and generation of in vivo phase diagrams

Cells were identified, segmented, and their fluorescent signal (median, average, minimum, maximum, 10th, 20th, …, 90th percentile fluorescence) as well as additional cell properties were identified using custom algorithms^61^ in FIJI^62^, and exported as tabulated files. Condensates were identified in each cell independently, in a multistep process: (i) we calculated the median fluorescence intensity of pixels in a given cell. (ii) we identified the largest region composed of pixels with an intensity 3-fold above the median. If such a region existed, showed a circularity above 0.4 and an area above 9 pixels, the cell was deemed to contain a condensate.

Tabulated data resulting from image analyses were loaded and analyzed with custom scripts in R. To convert fluorescent intensities to cytosolic concentrations, His-tagged Venus and FusionRed were purified using the GE Healthcare His GraviTrap system. Serial dilutions of each protein were generated, fluorescence intensities were recorded, and a linear model was fitted (Figure S3). A strain expressing only the two fluorescent proteins served to calibrate fluorescence signals. This strain was included in the experiments and analyzed alongside the serial dilution. Fluorescent signals of the experiments were normalized according to the normalizing strain and cytosolic concentrations were inferred from the regression of the purified proteins. Finally, cells with condensates were excluded, and the median cytosolic concentrations of YFP and RFP were plotted against each other.

### Fluorescence recovery after photobleaching (FRAP)

A macro created in VisiView 2.0 ® software was written to capture images on the red channel in rapid succession during the course of a FRAP experiment. Photobleaching was achieved with a 405 nm laser pulse lasting 20 ms after the 10th frame of the acquired series. The RFP channel exposure was set to 50 ms. Images were acquired every 100 ms. 250 frames for a total acquisition time of 25 seconds were acquired.

### FRAP data analysis

Custom macros were created in FIJI^62^ to extract quantitative data from the image series. Data was extracted from the non-bleached area and the bleached area by first manually selecting two pixel coordinates, firts at the center of the bleached region and second at the center of the non-bleached region. Then, a circular region of interest (ROI) of 6 pixels in diameter was generated. Since small movements of the condensate can occur during the movie, we generated 42 additional adjacent ROIs by translation of either 0.5, 1, 1.5 or 2 pixels in all directions, generating 6, 8, 12, or 16 ROIs for each distance respectively. Then, the average intensity of each ROI was extracted for every frame of the image series. The ROI intensities were subsequently analyzed with custom scripts in R. First, for each of the two locations (bleached and unbleached), we averaged 5 sub-ROIs showing either the lowest (bleached area) or highest total fluorescence intensity (non-bleached area). For each frame, the intensity recorded for the bleached area was divided by the intensity of the non-bleached area. Finally, the values were normalized as follows: 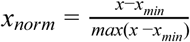, where x is the ratio of integrated pixel intensities measured in the bleached over unbleached ROI, and x_min_ is the minimum value of x across the image series.

## Acknowledgments

We thank Jeffrey Gerst and Rohini-Ravindran Nair for sharing plasmids of the MS2 system, Schraga Schwartz for the bRA89 plasmid, Francesco Sciortino for helpful discussions and suggestions and Harry Greenblatt for help with computer systems. This work was supported by the Israel Science Foundation (2179/14), by the European Research Council (ERC) under the European Union’s Horizon 2020 research and innovation programme (grant agreement No. 819318), by a research grant from A.-M. Boucher, by research grants from the Estelle Funk Foundation, the Estate of Fannie Sherr, the Estate of Albert Delighter, the Merle S. Cahn Foundation, Mrs. Mildred S. Gosden, the Estate of Elizabeth Wachsman, the Arnold Bortman Family Foundation. E.D.L. is incumbent of the Recanati Career Development Chair of Cancer Research. L.R. acknowledges support from the European Commission (Marie Skłodowska-Curie Fellowship No. 702298-DELTAS). S.A.S. thanks the BSF and the ISF program and acknowledges the historical generosity of the Perlmann family foundation. S.K.N. acknowledges support from the Koshland foundation.

## Authors contributions

M.H., J.M.G, and E.D.L. designed the research and synthetic protein system -- M.H., J.M.G. performed the experiments with help from Y.N. -- E.L., L.R. and J.K.P.D developed the theoretical framework for modeling the system based on patchy particles -- S.N. and S.S. developed the theoretical framework for modeling the system based on a lattice model -- A.S. wrote the image analysis scripts; E.S. carried out electron microscopy experiments -- M.H. and E.D.L. wrote the manuscript with input from all authors.

