## Supplementary material for "Designer protein assemblies with tunable phase diagrams in living cells": Supplemntary Material

**Table S1. Properties of the components of our synthetic minimal system.**

| Building Block | Valency | ORF | Oligomerization Domain | Dist. between oligomer termini | Linker 1 | Interaction Domain | Linker 2 | Fluorophore | Size (kDa) | Iso-electric Point |
|---|---|---|---|---|---|---|---|---|---|---|
| 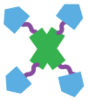 | 4 | Venus-E9-p53 | 1AIE (*1*) | ~4nm | GGSGS | **E9** | GGSGG | Venus | 46.22 | 6.69 |
| 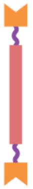 | 2 | NES-Im2-4LTB-FusionRed | 4LTB (*2*) | ~18nm | GGSGGS | **Im2 :** WT D33L E30A P56A V37A | GSGSG | Fusion Red | 60.72 | 5.02 |

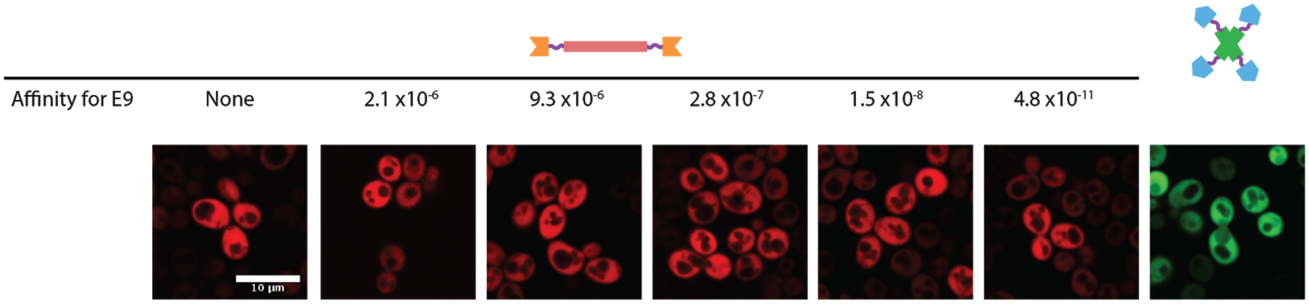


**Figure S1. The components do not form condensates when expressed individually.** Haploid cells expressing only one of the building blocks show a homogenous distribution of fluorescence throughout the cytoplasm. The left-most image shows cells expressing the dimer component lacking the Im2 domain. The next images show cells expressing the variants of the dimer component in absence of tetramer component. The right-most image shows cells expressing the tetramer component in the absence of the dimer component.

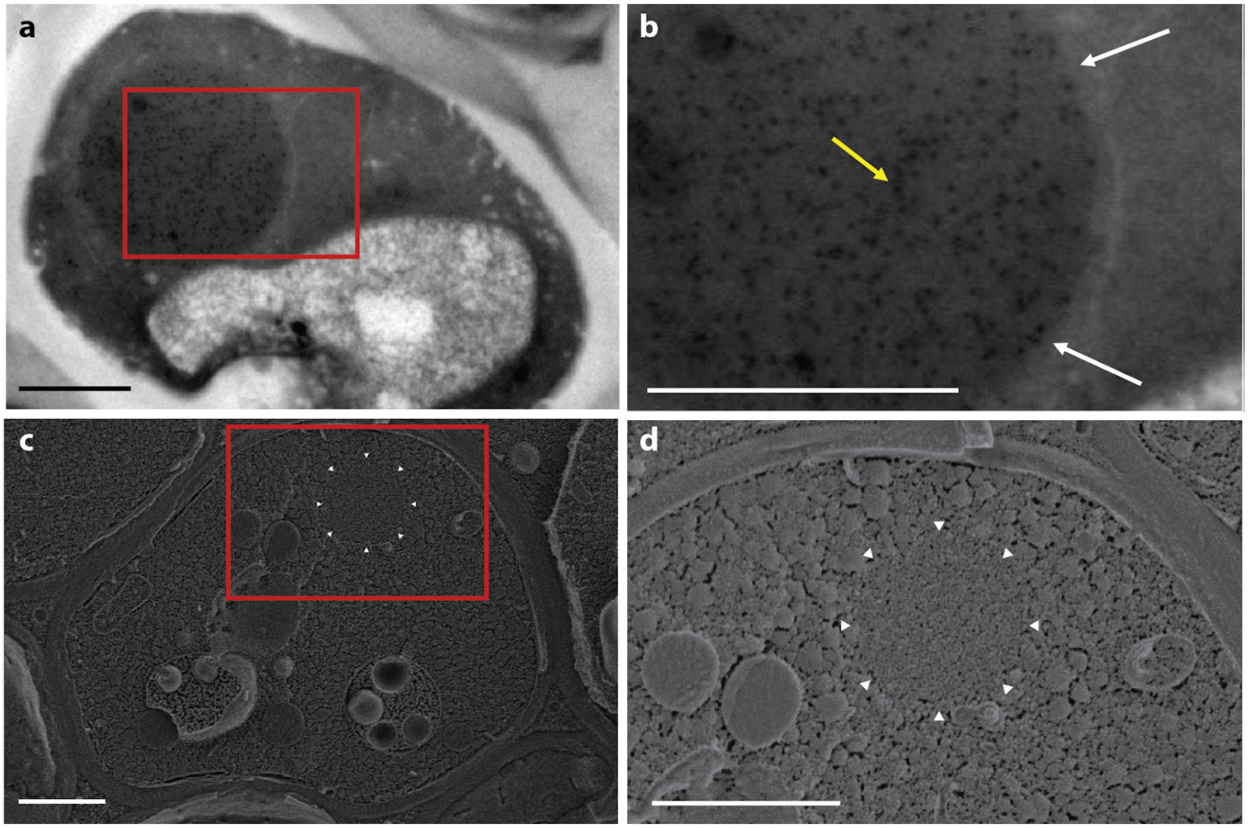


**Figure S2. The synthetic condensates are not membrane bound. a**. Transmission electron microscopy (TEM) micrograph of fixed and sectioned yeast shows a condensate formed by our minimal system, in the cytoplasm. **b.** The yellow arrow points to one of several 10 nm gold-labeled anti-GFP antibodies, confirming the identity of the designed compartments. White arrows highlight lack of membrane surrounding the compartment. **c.** Scanning electron microscopy micrograph of cells frozen at high-pressure and cryo-fractured reveals the mosaic of amorphic cytoplasm. The region outlined by white carets exhibits a distinct ultrastructure **d**. Increased magnification of a suspected condensate within the cytoplasm, outlined with white carets. This ultrastructure has no visible membrane. Scalebar 1 μm.

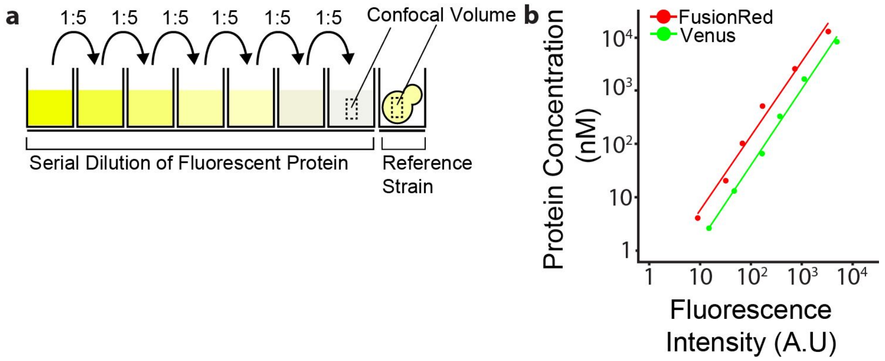


**Figure S3. Conversion of fluorescent intensities to concentrations. a**. The fluorescent proteins Venus and FusionRed harboring a HIS-tag were purified and prepared by serial dilution to span concentrations ranging from 41μM (Venus) or 64 μM (FusionRed) to 3nM (Venus) or 4nM (FusionRed). A negative control (without fluorescent protein) was included in the series. Mean fluorescence intensities for this serial dilution were measured alongside the normalizing strains' fluorescence intensity. **b.** A linear regression was used to determine the fluorescence units per protein concentration.

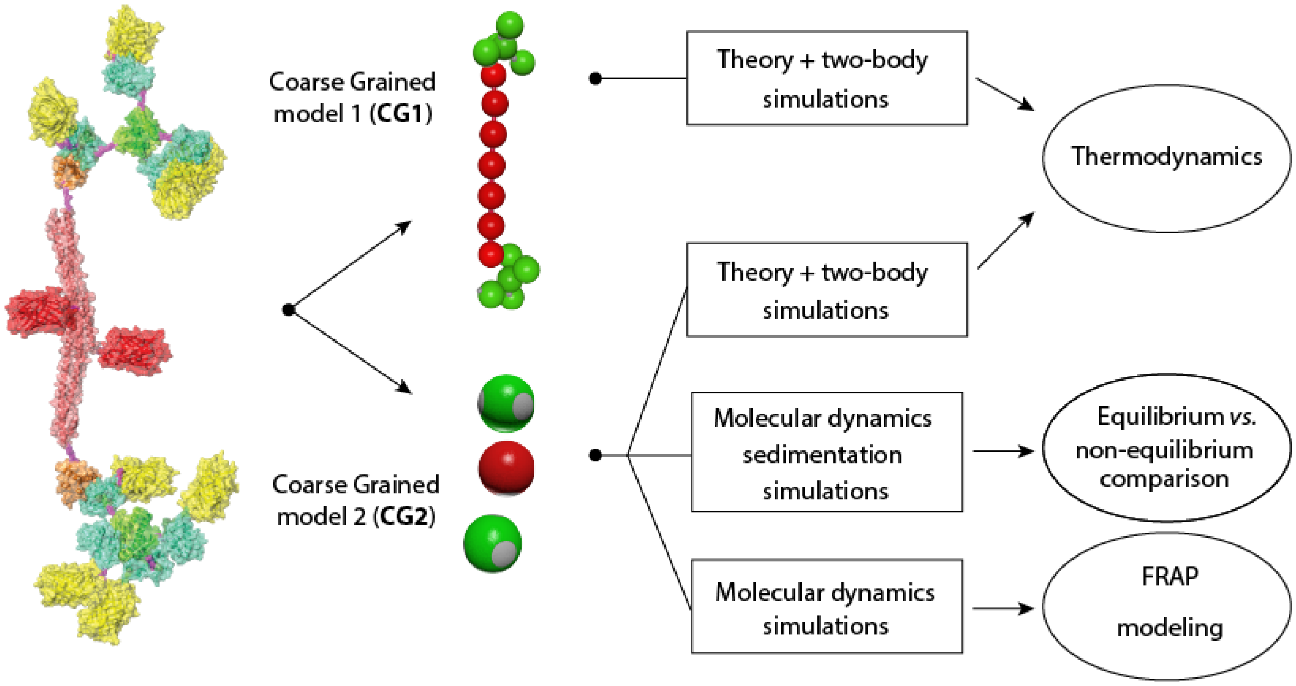


**Figure S4. Visual summary of the patchy particle models and of the analyses associated with each model.** A first coarse grained model (CG1) captures the geometry of the protein components: a long dimer (red) and small tetramer (green). The second model (CG2) uses one spherical particle per component and only reflects their valency, with four and two interaction patches for the tetramer and dimer respectively. We used a thermodynamic framework (Text S2.1) to describe the one- or two-phase equilibrium of both coarse grained models CG1 (Text S2.2) and CG2 (Text S2.3). Further, we used the model CG2 in molecular dynamics sedimentation simulations to investigate how the phase diagram of our system changes when it is out of equilibrium (Text S2.4). Finally, we employed model CG2 to model the effects of (i) density and (ii) fraction of bonded sites, on the diffusivity of components within the condensates (Text S3).

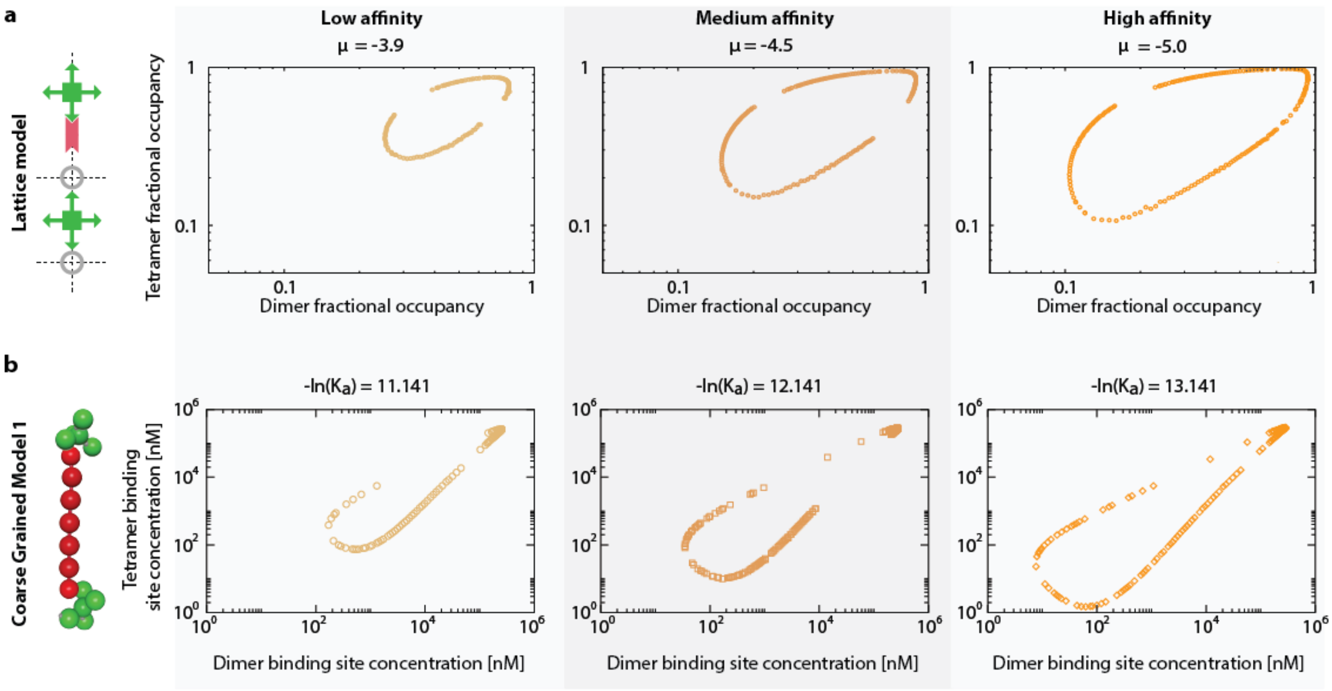


**Figure S5. Impact of affinity on the phase diagram of the dimer-tetramer system. a.** We used a lattice model ( Text S1) of the dimer-tetramer system. In the square lattice, concentration is measured by fractional occupancy of edges and vertices by dimers and tetramers respectively. We calculated the binodal of this system in the plane corresponding to the fractional occupancy of dimer (x-axis) and tetramer (y-axis). Affinity increases in panels from left to right, where μ is the binding energy in units of kT of a linker and one arm of the tetravalent molecule. Higher affinity (larger μ) increases the fraction of the phase separated region. **b.** We used mean-field theoretical calculations of patchy particles matching the geometry of the proteins. The binodal is calculated in the plane corresponding to the concentration of dimers (x-axis) and tetramers (y-axis). Affinity (which is linked to the energy and entropy associated with the formation of a bond, see Text S2.1) increases from left to right.

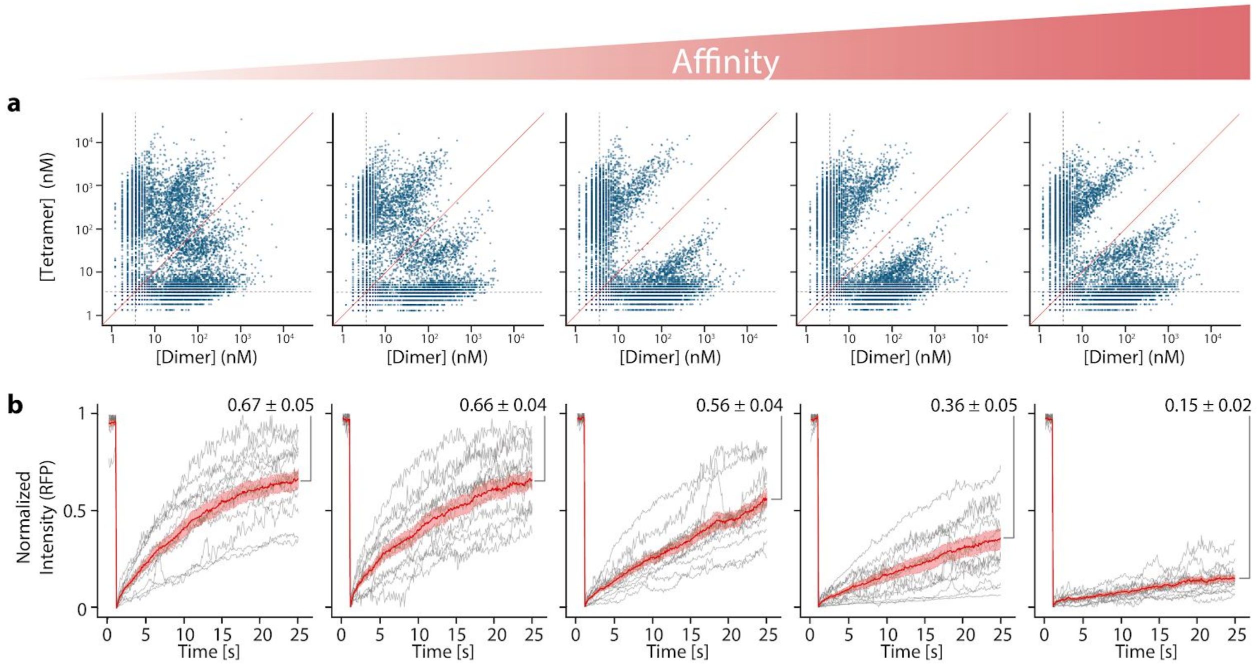


**Figure S6. *In vivo* phase diagrams and fluorescence recovery profiles observed with different affinities. a**. *In vivo* phase diagrams observed for all five affinities. Concentrations correspond to those of the binding sites (not of the dimer and tetramer complexes). The red line highlights the diagonal, where the concentrations of binding sites of dimer and tetramers is equal. The grey dotted lines show the lower limit of concentrations that can be reliably estimated. **b**. Fluorescence recovery profiles of photobleached condensates for different interaction affinities between the components. Grey lines show individual experiments and the red line corresponds to the mean recovery observed across ten independent condensates. The transparent red area indicates the standard error. The mean recovery after 25 seconds and associated standard error are given for each affinity.

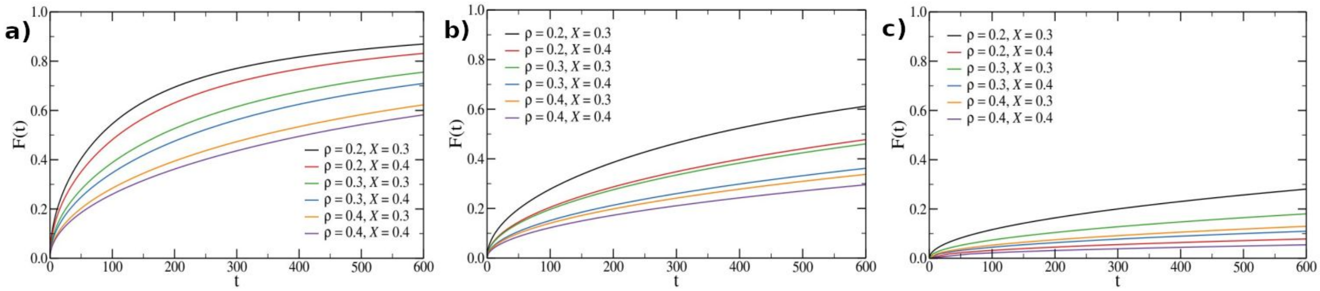


**Figure S7.** Simulated FRAP curves as a function of time for several values of density ( $\rho$ ), composition ($X$) and three different interaction strengths: a) $\frac{\varepsilon}{k_B T} = 8$ , b) $\frac{\varepsilon}{k_B T} = 10$ , c) $\frac{\varepsilon}{k_B T} = 50$ . Simulation were carried out as described in Text S3.

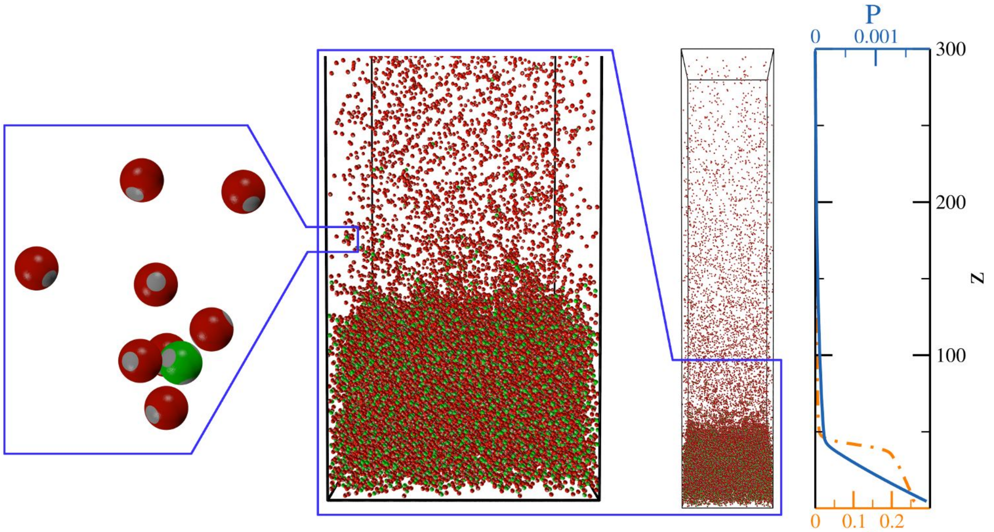


**Figure S8.** Snapshot of a system with $x_i = 0.25$. In the central panel, a blow up of the region close to the interface is shown; on its right, the entirety of the system is visible. On the left panel, a few representatives of divalent (red) and tetravalent (green) particles are shown, as a further blow up. On the right panel, the density (dashed and dotted line) and the pressure (solid line) are reported as a function of the height z for the same system.

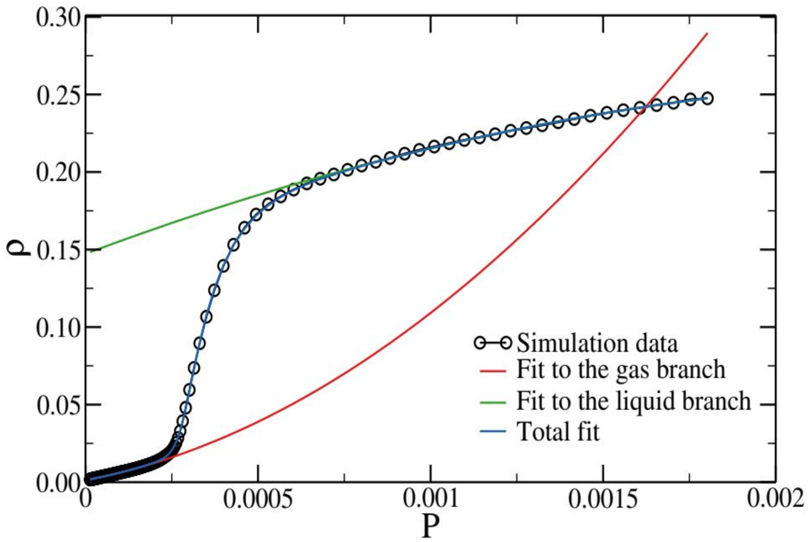


**Figure S9.** The function $\rho(P)$ for the equilibrium system with $x_i = 0.25$. The continuous curves are fits to the single branches (Eq. (6), red and green curves) and to the whole simulation data (Eq. (7) blue curve).

### Text S1. Physical basis of the lattice model of tetramers, dimers and solvent

We consider a lattice model and denote the tetramers as A molecules that can occupy the vertices and the dimers as B molecules that occupy the bonds of the lattice. Solvent molecules S, can occupy either the vertices or bonds. This allows us to focus on the bonding of the nearest-neighbor dimers and tetramers in a way that accounts for the valency of the tetramers. The physical origin of the phase separation lies in the tetramer-dimer attraction, which is the only interaction energy in this simplified model; nearest-neighbor tetramers or dimers separated by solvent molecules do not interact. Higher order neighbor interactions are neglected and the zero of energy is set by the tetramer-solvent and dimer-solvent interactions, which we take to be equal for simplicity. A schematic depiction of the system is shown in Fig. 2b.

The physical origin of phase separation of molecules in solution is the attraction between them, which, in the appropriate concentration, temperature and interaction-strength range, dominates the entropy of mixing. In our system of dimers and tetramers, there is an indirect, tetramer-tetramer attraction originating in their binding to mutual dimers. The essence of this effect can be seen by considering two tetramers and one dimer. It the two tetramers are relatively far, separated by the aqueous phase (Fig. 2b, grey inset), the dimer can bind to only one of them. However, if the two tetramers are in close proximity, the linker can bind to both, with a doubled binding energy; this results in an effective attraction of tetramers.

In equilibrium, this effect (for strong enough binding relative to the temperature) gives rise to a phase with a relatively high concentration of tetramers and dimers – with relatively high enthalpy, that coexists with a dilute phase – with a relatively high entropy. The thermodynamic criterion for coexistence is an equal chemical potential for each of the species (tetramer, dimer and solvent molecules) in the two phases. Our quantitative treatment captures these effects to predict the concentration, temperature and linker binding strength regimes where phase separation occurs. The statistical mechanics of this model are subtle due to the multivalency of tetramers and the ability of dimers to bind to only one tetramer arm. A mean-field theory and calculation that results in the phase diagrams shown in the main text is described elsewhere (to be published).

The experimental data corresponding to the interaction 1.5 x $10^{-8}$ M is about 18 $k_BT$ (Fig. 2g). The lattice model involves solving four nonlinear algebraic equations to find the equilibrium concentrations of the complexes and then using interpolation we find the analytical expression for the free energy that we finally use to find the binodal phase diagram numerically (to be published). This procedure makes it hard to numerically find the binodal for very large interaction strengths. The theory (to be published) shows that the minima of the phase diagrams vary exponentially with the interaction strength. For these reasons, we show an overlay of the theoretical binodal (and not a fit) on the experimental data. The agreement suggests that the shape of the phase diagrams obtained in the experiment and from the lattice model are similar.

### Text S2. Physical basis of the Simulations of Dimer and Tetramer Patchy Particles

Here we introduce an alternative approach to the lattice model, based on numerical simulations and mean-field theoretical calculations that qualitatively reproduces the behaviour of the experimental dimer-tetramer system. In order to simplify both theoretical and numerical descriptions, we take into account the effect of the solvent only implicitly. Thus, the system is composed of only two species of particles that are the counterparts of the synthetic proteins used in the experiments: the tetramers denoted as A molecules can form up to four bonds, and the dimers denoted as B molecules can form up to two bonds. In addition, only unlike particles can be bonded (i.e., a dimer can only bind to a tetramer).

The two coarse-grained (CG) models that we use are illustrated in Fig. S4. The first, Model CG1, attempts to capture the size, shape and flexibility of tetramers and dimers relatively realistically. The computed properties of these species can then be used as inputs into a theoretical approach (described in detail in Section 2.1) to derive the phase diagram. However, model CG1 is too complex for direct simulations of the phase separation behaviour to be feasible on reasonable time scales. We have therefore introduced a more coarse-grained model, Model CG2, in which the tetramers and dimers are represented by simple patchy particles. Although this model is less geometrically realistic of the experimental system, it is expected to capture the basic physics underlying the phase separation well and to exhibit a similar phase behaviour. As with Model CG1, we can use the theory of Section 2.1 to predict the equilibrium phase behaviour. However, we can also perform direct simulations of the phase separation behaviour. In particular, we have used the latter to analyze non-equilibrium effects on the observed phase separation at high affinities.

#### 2.1. Thermodynamics used in both coarse-grained models (CG1 and CG2).

To compute the phase diagram for Models CG1 and CG2, we use a theoretical approach that has been shown to work well for DNA bulk fluids(*3*, *4*). The first task is to compute the free energy of the system at a given state point (e.g. given temperature, overall concentration and composition). The expression for the free energy takes into account the overall translational entropy, the entropy of mixing of the two species, the effects of excluded volume, and bonding (where the number of bonds is given by a simple chemical equilibrium). Then, the condition that the chemical potentials (for both species) are equal in the coexisting low-density and high-density phases is used to map out the phase boundaries. The expressions for the free energy require a number of model-specific inputs that characterize the excluded volume and the free energy of a single bond. These are obtained from simulations of the relevant species. While the theoretical framework and how it is used to compute the phase diagrams has been described previously(3), we provide the reader with a self-consistent description specific to the present work.

We calculate the Helmholtz free energy per particle of the binary mixture following Wertheim(5, 6):

$$\beta f_{tot} = \beta f_{ref} + \beta f_{bond}$$

where $\beta = \frac{1}{k_B T}$ is the absolute inverse temperature and $f_{ref}$ is a reference free energy, for which we use the following expression that takes into account the ideal gas, mixing and excluded volume contributions:

$$\beta f_{ref} = log\,(\rho v_0) - 1 + x log\,(x) + (1-x)\,log\,(1-x) + \beta f_{ex}$$

$v_0$ is a reference volume, $x = \frac{N_A}{(N_B+N_A)}$ is the composition ($N_B$ and $N_A$ are the number of particles of type $B$ and $A$, respectively) and $\rho$ is the overall number density. The last term is the excess free energy, which we choose to represent using the second virial approximation:

$$\beta f_{ex} = \rho(x^2 B_{AA} + 2x(1-x)B_{AB} + (1-x)^2 B_{BB})$$

where $B_{AA}$ and $B_{BB}$ are the second virial coefficient of the first and second species and $B_{AB}$ is the inter-species one. The values of $B_{AA}$, $B_{BB}$ and $B_{AB}$ have been computed using the relation

$$B_2^{ij}(T) = -\frac{1}{2}\int_0^\infty 4\pi r^2 \left(exp\left(-\frac{V_{ij}(r)}{k_B T}\right) - 1\right) dr \tag{1}$$

where $V_{ij}(r)$ is the effective intra-molecular pair potential of the reference system, here taken as a system where attraction, and hence inter-particle bonding, is disabled. These values are computed through dedicated numerical simulations (see Sections 2.2 and 2.4).

Following Wertheim , the bonding free energy is given by

$$\beta f_{bond} = x\left[M_A log\,(X_A) - \frac{M_A X_A}{2} + \frac{M_A}{2}\right] + (1-x)\left[M_B log\,(X_B) - \frac{M_B X_B}{2} + \frac{M_B}{2}\right]$$

$M_A$ ($M_B$) being the valence of particles of type $A$ ($B$), $p_A = 1 - X_A$ and $p_b = 1 - X_B$ being the probabilities for any patch of type $A$ and $B$ to be part of a bond. Following previous work(7), these probabilities are related to the number of bonds $n_b$ and to the number of particles of the corresponding species $N_A$ ($N_B$) as:

$$p_A = \frac{n_b}{M_A N_A}; p_B = \frac{n_b}{M_B N_B} = \frac{p_A M_A N_A}{M_B N_B} \tag{2}$$

Furthermore, $X_A$ ($X_B$) can be expressed in terms of the density $\rho$ and the composition $x$ of the mixture using the law of mass action

$$\frac{n_b}{(M_A N_A - n_b)(M_B N_B - n_b)} = \frac{v_0}{V} e^{-\beta F_b} = \Delta \tag{3}$$

where $v_0$ is a reference volume and

$$e^{-\beta F_b} = \frac{v_b(\rho)}{v_0}\left[e^{\beta\epsilon} - 1\right]$$

where $v_b$ is to be interpreted as a bonding volume and $\epsilon$ is the energy associated with the formation of a bond. We approximate $\Delta$ with the following relation:

$$\Delta = 4\pi \int_{\sigma}^{\sigma+\delta} g_{HS}(r) \langle f(12) \rangle_{\omega_1,\omega_2} r^2 dr \tag{4}$$

where $g_{HS}(r)$ is the reference hard-sphere fluid pair correlation function, $\langle f(12) \rangle_{\omega_1,\omega_2}$ represents an angle-average of the Mayer function over all orientations of particles $1$ and $2$ at fixed relative distance $r$.

Notice that the law of mass action can be re-written in terms of the affinity $\frac{1}{K_a}$ , *i.e.* $\frac{1}{K_a} = v_b(\rho)\, exp(\beta\epsilon)$. In this way, one can directly connect theory and experiments, provided that one measures the bonding volume in the correct units. Combining Eqs. (2) and (3), we obtain the following expressions for $X_A$ and $X_B$

$$X_A = \frac{-(u_A+\Gamma^{-1})+\sqrt{(u_A+\Gamma^{-1})^2+4\Gamma^{-1}}}{2}$$

$$X_B = 1-(1-X_A)\,\frac{M_A x}{M_B(1-x)}$$

where $\Gamma = M_A x \rho v_b (e^{\beta\epsilon}-1)$ and $u_A = \frac{M_B}{M_A}\frac{1-x}{x}-1$. The approach we use to find the coexistence region requires the computation of the Gibbs free energy per particle, $g(P,T,x)$. We first formally write

$$g^*(T,P,\rho,x) = f_{tot}(x,T,\rho) + \frac{P}{\rho},$$

then we minimize $g^*$ as function of $\rho$ at fixed temperature, composition and pressure to find $g(P,T,x)$, as we cannot invert the $P(\rho)$ relation. Minimization is carried out using a standard minimization algorithm(*3*). Phase coexistence requires the equality of the chemical potentials between the two phases for both species at the same time. Coexistence points at fixed temperature and pressure are thus found imposing

$$(P,T,x_1) = \mu_A^2(P,T,x_2),\ \mu_B^1(P,T,x_1) = \mu_B^2(P,T,x_2) \tag{5}$$

where $\mu_k^j$ is the chemical potential of the species $k = A,B$ in the phase $j$. Each solution ($x_1,x_2$) of the above system of equations defines a pair of concentrations ($\rho_A^1$ ,$\rho_B^1$ ) corresponding to the dilute phase and another pair of concentrations corresponding to the dense phase ($\rho_A^2$ ,$\rho_B^2$ ). In order to solve Eqs. (5), we use a nonlinear solver (modified Powell algorithm)(*3*). We remark that solving the system of equations Eqs. (5) is a way to implement the *common tangent construction* for a binary mixture.

### 2.2. Description of the first coarse grained model (CG1).

This coarse-grained model attempts to provide a realistic description of the experimental system's geometry whilst being simple enough to allow the model-specific inputs required by the theory of Section 2.1 (namely the second virial coefficients $B_{AA}$, $B_{BB}$, $B_{AB}$ and the bonding free-energy $\Delta$) to be computed in simulations of the model. The tetramers are represented by 5 beads. The central particle represents the tetrameric p53 complex, and is connected by flexible linkers to four identical particles that represent the E9 proteins. By contrast, the dimers are made by 7 beads. The five innermost particles represent the 4LTB complex, whereas the two outer beads represent the Im2 proteins.

Here we detail the model we use to compute the second virial coefficients $B_{AA}$, $B_{BB}$, $B_{AB}$ and the bonding free-energy $\Delta$ that are used as inputs in the theory.

Tetramers are composed of a central bead and four identical beads that are connected to the former and bear a single attraction point (a patch). All these beads have diameter $\sigma_l$. By contrast, a dimer is modelled as a chain whose middle part is made by five beads of diameter $\sigma_s = 1.15\,\sigma_l$ connected to two outer beads of diameter $\sigma_l$ bearing a patch each. In the following, energies are measured in units of $k_B T$ and lengths in units of $\sigma_l$.

Any two spheres $i$ and $j$ feel a mutual repulsion given by

$$V_{CM}(ij) = A\left(\frac{\sigma_{ij}}{r_{ij}}\right)^{200}$$

where $A = 1$ and $\sigma_{ij} = \frac{(\sigma_i + \sigma_j)}{2}$.

The surface of the (four for the tetramers and two for the dimers) outermost particles is decorated with a single patch. The attractive interaction between two patches on particles $i$ and $j$ is given by

$$V_P(ij) = -\epsilon exp\left[-\frac{1}{2}\left(\frac{r_P^{ij}}{0.12\sigma_l}\right)^{10}\right]$$

where $r_P^{ij}$ is the distance between two spherical patches and $\varepsilon = 1.001$.

There are sets of springs acting between the centre of masses of all particles. In particular, in the tetramer there is a spring connecting each patchy bead with the corresponding central bead, whereas in the dimer all particles in the chain are connected to their neighbour(s). All these springs are modelled with a FENE potential of the form

$$V_F(r) = -15A\frac{r_F^2}{\sigma_l^2}log\left(1 - \frac{r^2}{r_F^2}\right)$$

where $r_F = 2.25$. Note that we always use $\sigma_l$ in the denominator of the prefactor, irrespective of the species of the two particles.

The dimers are kept stiff by a three-body interaction of the form

$$V_{3B}(\theta) = -k(1 - cos\theta)$$

where $k$ controls the bending stiffness and $\theta$ is the angle formed by three consecutive monomers. We set high stiffness with $k = 5$ for the central beads and a relaxed stiffness with $k = 0.5$ for the outermost triplets, to account for the flexibility introduced by the linkers in the experimental system. The same three-body potential with relaxed stiffness (k = 0.5) is also applied to the beads in the tetramers (where we consider all patchy-centre-patchy triplets) so as to favour tetrahedral symmetry. The relative average sizes of the two constructs are compatible with the experimental ones. A dimer (in red) linked to two tetramers (in green) are depicted in Fig. S4, with interaction patches shown in grey.

For this model the computation of the second virial coefficients yields $B_{AA} = 23.0158\sigma_l^3$, $B_{AB} = 38.6495\sigma_l^3$, $B_{BB} = 64.8598\sigma_l^3$. To compute the bonding free energy we make the approximation $g_{HS}(r) = 1$, valid at low densities, and we estimate the angle-average of the Mayer function using numerical simulations with a generalized Widom insertion method(*8*), finding $v_b(\rho) = v_b = 0.02109\sigma_l^3$. By inspecting the x-ray crystallographic structure of the tetrameric protein domain (PDB code: 1AIE) we find $\sigma_l$ = 3.8 nm, which allows to convert simulation units to real ones: the conversion factor between number density in simulation units and nanomolar is given by $(\sigma_l \cdot N_A)^{-1}$ = 3.3044E7. With this model, the average distance between two tetramers linked by a dimer is $d_{CG1} \approx 31.3$ nm.

### 2.3. Theoretical coexistence regions calculated for the CG1 model

We report here the coexistence regions as computed using the theoretical framework described above. The values of the second virial coefficients are fixed to the ones reported in Section 2.1. We define as $\rho_A$ and $\rho_B$ the concentrations of monomeric units forming the dimers and tetramers, respectively, so $\rho_A = 2[A]$ and $\rho_B = 4[B]$, where [A] and [B] are the concentrations of dimers and tetramers, respectively.

We report the coexistence regions in the plane $\rho_A - \rho_B$ for different values of the affinity $-ln(K_a)$, defined as $-ln(K_a) = ln(v_b) + (\beta\epsilon)$ (Fig. S5); in practice, we keep $v_b$ fixed and we change $\beta\epsilon$. We observe that the coexistence regions enlarge and expand to more extreme values of $\rho_A$ and $\rho_B$ upon increasing the affinity, as observed in the experiments.

### 2.4. Description of the second coarse grained model (CG2) and out-of-equilibrium simulations

The CG1 Model is computationally too expensive to directly simulate phase separation, so we introduce the CG2 Model. This model is a binary mixture of simple patchy particles, where the tetramers are represented by a single particle with a tetrahedral arrangement of patches and the dimers by a single particle with two patches that are diametrically opposite. There is an attractive interaction between the patches on the tetramers and dimers, where $\varepsilon$ is the patch-patch attraction strength. The details of the model can be found in Ref.(*9*). In order to map the model's internal unit of length, set by the particle diameter $\sigma$ , to the experimental one we use the average distance between two bonded tetravalent particles. This quantity, which is essentially temperature-independent, is equal to $d_{CG2} \approx 2.25\,\sigma$. Setting $d_{CG1} = d_{CG2}$ yields $\sigma = 13.9$ nm.

In order to use the theoretical approach of Section 2.1 to predict the phase diagram for the patchy particle model, we first need to obtain the model-specific inputs required by the theory. First of all, we have computed the equation of state from which we can directly estimate the excluded-volume contribution to the reference free energy $f_{ex}$ (rather than use Eq.(1)). Interestingly, we see that the numerical data can be quantitatively described by the Carnahan-Starling excess free energy using an effective diameter $\sigma = 1.1$ and, consequently, we have decided to use the standard reference hard-sphere fluid pair correlation function $g_{HS}(r)$ in Eq. (4). By means of dedicated simulations and standard fitting procedure, we obtain the following functional form for $\Delta$

$$\Delta(\rho, \varepsilon) = (0.016166 + 0.026265\rho + 0.0017927\rho^2 + 0.06624\rho^3 - 0.014791\rho^4)\, exp(\beta\epsilon).$$

The CG2 model is also used to model FRAP traces ( Text S3).

Here we numerically estimate the phase diagram of a binary mixture of divalent and tetravalent particles in order to help understand the origin of the asymmetry of the phase diagram of the highest-affinity experimental system. To this end, we run simulations at an attraction strength at which no bonds can break over the course of a simulation ( $\frac{\varepsilon}{k_B T} = 50$ ). We run two sets of simulations: in one we use a regular patchy-particle model which, for this value of the attraction strength, is out of equilibrium within the observation time frame considered, whereas in the other we complement the model with a swap mechanism that allows the system to reach equilibrium even in the $T \to 0$, or, equivalently, $\frac{\varepsilon}{k_B T} \to \infty$, limit(*9*). We evaluate the phase behaviour by performing sedimentation simulations: we put $N = N_4 + N_2 = 36000$ particles in a box of size $L \times L \times L_z$, where $L_z = 400$ and $L = 60$, and apply a constant force $-g\hat{z}$ (where $g = 0.0002$ in simulation units) on all particles. We run several simulations where we vary the overall mole fraction of tetramers $x_i \equiv \frac{N_4}{(N_2+N_4)}$ .

An example of a configuration where the sedimentation equilibrium has been reached is shown in Fig. S8, together with a blow up of the interface and of a small portion of the system that shows a detailed view of single particles.

The main outputs of this procedure are the density, tetramer mole fraction and pressure profiles ( $\rho(z)$, $x(z)$, and $P(z)$, respectively). We can then extract $P(\rho)$, $P(x)$, $x(\rho)$, *etc.* by plotting these quantities parametrically (eliminating $z$). The function $\rho(P)$ is particularly interesting and will be the focus of the remaining part of this section.

The black dots in Fig. S9 shows $\rho(P)$ for the case $x_i = 0.25$ (in equilibrium). The jump in the density signals the occurrence of a phase separation which, in the thermodynamic limit $L, N \to \infty$, would be discontinuous. We build on Ref.(*10*) and estimate the coexisting $\rho$ and $x$ by fitting the numerical curve with a function that interpolates between the gas and the liquid branches. We first identify the gas and liquid branch by eye and fit the two with the (phenomenological) functional form

$$\rho_i(P) = A_i P + B_i P^2 + C_i \qquad (6)$$

where $i$ is either $g$ (for gas) and $l$ (for liquid) and $A_i$, $B_i$ and $C_i$ are three fitting parameters for each branch. Once the equations of state for the two branches are known, we interpolate between them with the function

$$\rho(P) = \rho_l + S(P)\left(\rho_g - \rho_l\right) \quad (7)$$

where $S(P)$ is a skewed sigmoidal function given by

$$S(P) = \frac{1}{\left(1+exp\left(\frac{P-P_c}{D}\right)\right)^E}$$

where $P_c$, $D$ and $E$ are three fitting parameters. The only parameter with a physical meaning is $P_c$, which is the pressure at coexistence. Fig. S9 shows an example of the resulting curves. Once $P_c$ is known, the densities of the coexisting phases are computed as $\rho_g^{coex} \equiv \rho_g(P_c)$ and $\rho_l^{coex} \equiv \rho_l(P_c)$. Knowing these values, the mole fractions of the tetramers at coexistence are estimated as $x_g^{coex} \equiv x\left(\rho_g^{coex}\right)$ and $x_l^{coex} \equiv x\left(\rho_l^{coex}\right)$. The partial densities are then obtained as $\rho_4 = \rho x$ and $\rho_2 = \rho(1-x)$. We run this analysis for systems with $x_i$ ranging from $0.2$ to $0.45$.

The set of coexisting phases that we observe for the equilibrium and non-equilibrium systems are plotted in Fig. 3e of the main text. The non-equilibrium apparent phase diagram is smaller than its equilibrium counterpart. The difference is most evident at high density (not accessed in experiments) and in the left lower quadrant, but not in the left upper quadrant, in agreement with experiments (see Fig. 3a of the main text). The source of the asymmetry between the two branches is not yet fully understood, but the agreement between simulations and experiments seems to indicate a physical origin that is present even in the simple patchy model we use.

The theoretical phase diagram reported in Fig. 3e of the main text has been obtained using the theoretical approach presented in Section 1.1 along with the model-specific inputs detailed in Section 2.1.

### Text S3. Modeling the FRAP data

Following Soumpassis(*11*), the normalised fluorescence recovery function can be written as

$$F(t) = exp\left(-2\frac{\tau_D}{t}\right)\left(I_0\left(2\frac{\tau_D}{t}\right) + I_1\left(2\frac{\tau_D}{t}\right)\right),$$

where $\tau_D = \frac{r_2^2}{(4D)}$ is the time required for a protein to diffuse over a distance equal to the bleach radius $r_e$, $D$ is the diffusion constant and $I_0$ and $I_1$ are modified Bessel functions. We compute $D$ in simulations as

$$D = \lim_{t\to\infty} \frac{\langle r^2(t)\rangle}{6t}$$

where $\langle r^2(t)\rangle$ is the mean-squared displacement. We run constant-temperature simulations of the CG2 model introduced above, but without any gravitational attraction. We compute $D$ for different values of $\rho$, $x$ and interaction strengths $\frac{\varepsilon}{k_BT}$, where $\varepsilon$ is the patch-patch attraction strength. Fig. S9 shows $F(t)$ for three different interaction strengths ($\frac{\varepsilon}{k_BT}$ = 8, 10, 50) and for the same range of densities.

The numerical results are in qualitative agreement with experiments (see Fig. 3C of the main text), showing that, as the protein-protein interaction strength gets stronger and stronger, not only the recovery gets slower and slower, but also the spread of the curves (which in experiments can at least be partially traced back to the different concentration inside the *foci*) decreases.
